# Active and unusually expanded *PIF/Harbinger* transposable elements in the *Caenorhabditis inopinata* genome

**DOI:** 10.64898/2026.05.26.728016

**Authors:** Kazuki Sato, Xiaodan Jin, Shun Oomura, Kazuma Kawahara, Simo Sun, Akemi Yoshida, Nami Haruta, Asako Sugimoto, Taisei Kikuchi

## Abstract

**Background:** Transposable elements (TEs) serve as powerful drivers of genome innovation but also threaten genome integrity. The *PIF/Harbinger* superfamily is distinctive among DNA transposons because mobilisation typically requires proteins, a DDE transposase and a MADF DNA-binding protein. *Caenorhabditis inopinata*, the closest known relative of *C. elegans*, has a TE-rich genome and lacks multiple components of the ERGO-1–class endogenous small-RNA pathway, making it a useful system for examining TE dynamics in a distinct host context. We identified a spontaneous dumpy mutant of *C. inopinata* caused by insertion of a *PIF/Harbinger*-family element into the coding region of *Cin-dpy-11*. The inserted element, designated *Harbinger-1M_cIno*, belongs to the *Turmoil2* lineage originally defined in *C. elegans* and retains a MADF domain but lacks a recognisable DDE transposase ORF. Genome-wide curation recovered 258 related copies, revealing a strongly asymmetric family structure. Short noncoding derivatives were predominant, MADF-bearing derivatives were expanded and only one DDE-bearing locus retained an apparently intact transposase gene, suggesting that DDE and MADF functions are partitioned across distinct elements and may be supplied in trans during mobilisation. We also identified a second *PIF/Harbinger*-derived family, *Harbinger-2M_cIno*, associated with the *Turmoil1* lineage. This family comprises 1,376 copies and therefore records substantial past amplification, but it lacks a detectable DDE source, shows greater sequence divergence and more degraded terminal structures than *Harbinger-1M_cIno*. Together, these data indicate that the two *PIF/Harbinger* lineages in *C. inopinata* differ not in whether amplification occurred, but in when it occurred and whether present-day mobilisation competence has been retained.

## Introduction

Transposable elements (TEs) are ubiquitous in eukaryotic genomes and are major drivers of genome evolution (Bourque *et al*., 2018). While their mobilisation can compromise genome integrity and fitness, TEs also generate regulatory sequences, promote genome rearrangements, and contribute to coding and noncoding innovation.

Two major mechanistic classes of TEs are generally recognised. Class I elements (retrotransposons) mobilise via an RNA intermediate and are therefore often described as “copy-and-paste” elements. Class II elements (DNA transposons) mobilise as DNA, and many transpose through a canonical “cut-and-paste” mechanism. In canonical DNA transposons, the transposase recognises terminal inverted repeats (TIRs), excises the transposon sequence, and reinserts it elsewhere in the genome, typically generating short target-site duplications (TSDs) upon integration (Wicker *et al*., 2007). Transposable elements can broadly be classified as autonomous or non-autonomous according to whether they encode the machinery required for their own mobilisation. This distinction applies to both retrotransposons and DNA transposons, although the enzymatic machinery involved differs between them. Within DNA transposons, autonomous elements encode a transposase, whereas non-autonomous elements depend on trans-acting transposase supplied by related autonomous elements. Miniature inverted-repeat transposable elements (MITEs) are short, non-autonomous derivatives of TIR DNA transposons that lack a protein-coding open reading frame (ORF) but retain the terminal features required for mobilisation. They are generally thought to arise through internal deletion of autonomous progenitors, with abortive or interrupted gap repair representing one commonly proposed mechanism. Because direct sequence similarity to autonomous partners is often restricted to the terminal regions, the evolutionary origin of individual MITE families can be difficult to resolve. Once formed, however, such compact derivatives may undergo substantial copy-number amplification in the presence of a compatible trans-acting transposase (Rubin and Levy, 1997; Fattash *et al*., 2013).

The *PIF/Harbinger* superfamily is widely distributed across eukaryotes. *Harbinger* elements were initially identified in plant genome analyses, including *Arabidopsis thaliana* (Kapitonov and Jurka, 1999), and were later unified with maize *PIF* elements within a single *PIF/Harbinger* superfamily. In nematodes, related elements were first characterised in *C. elegans* as *Ce-PIF1* and *Ce-PIF2* and were subsequently linked to the *Turmoil* elements (Jurka and Kapitonov, 2001; Zhang *et al*., 2001).

*PIF/Harbinger* elements typically carry TIRs of highly variable length and generate 3-bp TSDs at a characteristic TWA target site upon insertion (Han *et al*., 2015). A characteristic feature of autonomous *PIF/Harbinger* elements is their two-gene organisation: one gene encodes a DDE transposase, typically with an associated helix-turn-helix-related DNA-binding region, and the other encodes a Myb/SANT-like DNA-binding protein, often referred to as a MADF- or trihelix-like factor (Wicker *et al*., 2007). Available genetic and heterologous reconstitution studies indicate that both proteins are required for efficient *PIF/Harbinger* transposition (Yang *et al*., 2007; Sinzelle *et al*., 2008; Hancock *et al*., 2010; Liu *et al*., 2024). In experimentally tractable systems, this Myb-like partner binds terminal or subterminal transposon sequences, interacts with the transposase, and promotes its accumulation in the nucleus, supporting assembly of a functional transposition complex at the element ends.

In animals, TE mobilisation is typically restrained by host defence systems, particularly small-RNA pathways (Siomi *et al*., 2011; Iwasaki *et al*., 2025). In *C. elegans*, piRNAs and endogenous siRNAs act together to suppress TE expression, largely through secondary 22G-RNA amplification and WAGO-mediated silencing, with the ERGO-1–class 26G pathway providing an additional upstream trigger on selected targets (Das *et al*., 2008; Guang *et al*., 2008; Buckley *et al*., 2012; Billi *et al*., 2014). By contrast, *C. inopinata*—the closest known relative of *C. elegans*—has a larger, TE-enriched genome and lacks several genes of the ERGO-1–class 26G-siRNA pathway, including *ergo-1, eri-6/7,* and *eri-9* (Kanzaki *et al*., 2018). Its genome also bears clear signatures of TE-associated restructuring, including disruption of the *her-1* orthologue by an LTR insertion (Kanzaki *et al*., 2018). Although these observations do not on their own establish globally elevated ongoing TE activity, recent work has demonstrated active *hAT*-family transposition in *C. inopinata*, including repeated mobilisation of Ci-hAT1 and expansion of related non-autonomous derivatives (Hatanaka *et al*., 2024).

*C. inopinata* therefore provides a valuable comparative framework for examining how divergence in small-RNA pathways and TE dynamics shapes genome structure and function in *Caenorhabditis*.

Here, we analysed a spontaneously arising dumpy (Dpy) mutant of *C. inopinata* and identified a novel insertion of a *PIF/Harbinger* DNA transposon, providing direct evidence of ongoing mobilisation. The newly inserted copy retains an intact MADF-like DNA-binding domain but lacks a DDE transposase ORF, indicating that it is a derivative in which only the MADF module has been retained. Genome-wide analysis revealed more than twenty highly similar copies of this MADF-only form, as well as a distinct locus encoding a DDE transposase but lacking a cognate MADF ORF, a genomic configuration consistent with mobilisation in trans. Phylogenetic and structural analyses place these MADF-only derivatives within the *Turmoil2* lineage. *Turmoil1*-related sequences are also present in the genome; however, no *Turmoil1*-type DDE transposase was detected, and the remaining MADF-containing loci have degraded TIRs, together suggesting that this lineage is no longer mobile. Thus, *C. inopinata* contains two *Turmoil*-related lineages with contrasting current states: a *Turmoil2* lineage that retains evidence of ongoing mobilisation and a *Turmoil1* lineage that appears to have degenerated. This contrast provides a tractable system for investigating how the modular MADF–DDE architecture influences lineage-specific TE evolution.

## Materials and Methods

### Nematode strains and maintenance

*C. inopinata* was maintained under modified *C. elegans* growth conditions as previously described (Kanzaki *et al*., 2018; Oomura *et al*., 2022). A spontaneous Dpy mutant, SA1635 [*tj217/tj217*], was isolated from laboratory cultures of the wild-type strain NKZ35 based on its shortened, stout body morphology at the L4 or adult stage. Because homozygous progeny from homozygous mothers were sterile, the allele was maintained in the heterozygous state by crossing to wild type. A wild-type population (SA1689), derived within several generations of the original isolate, was used as the control. Images were acquired directly from NGM plates with a Nikon SMZ1270 stereomicroscope and a Leica MC170 HD camera using LAS-EZ software.

### Whole-genome sequencing and variant calling

Genomic DNA was extracted from single adult males of SA1635 and SA1689, with five biological replicates per strain. DNA from individual worms was used to prepare sequencing libraries with the Illumina DNA Prep Kit (Illumina). Libraries were sequenced on an Illumina NovaSeq 6000 platform to generate 150-bp paired-end reads.

Reads were trimmed to remove adapter and low-quality sequences and mapped to the *C. inopinata* reference genome (v7.11) with SMALT v0.7.4 (https://www.sanger.ac.uk/resources/software/smalt/). Alignments were processed with SAMtools (Danecek *et al*., 2021) and Picard tools (http://broadinstitute.github.io/picard/). Small variants were called with GATK HaplotypeCaller (https://gatk.broadinstitute.org/hc/en-us) using filtering criteria of QUAL > 30 and depth ≥ 10×, and structural variants were identified using Manta v1.3.1 (Chen *et al*., 2016). Variants were annotated with ANNOVAR (Wang *et al*., 2010). Variants of interest were visualised in Artemis (Carver *et al*., 2012) to confirm breakpoint structures.

### PCR validation of the *Cin-dpy-11* insertion

Genomic DNA from SA1635 and SA1689 was amplified with primers located in intron 2 and the 3′ UTR of *Cin-dpy-11* (forward, 5′-gaagcagtcgtgaaaacaagagagg-3′; reverse, 5′-gcatatcacaggtacaatcccgtc-3′). The expected wild-type product size was 975 bp. When SA1635 genomic DNA was used as the template, an approximately 1-kb PCR artefact was also detected, likely because annealing between TIR sequences formed a hairpin-like secondary structure, resulting in skipping of the TE insertion. PCR products were analysed by Sanger sequencing. Sanger reads were aligned to the *Cin-dpy-11* genomic region to define the breakpoint structure and insertion coordinates by confirming TIR sequences at both ends.

### Repeat identification

RepeatMasker v4.1.8 was run with the parameters -a -s -e rmblast on *C. inopinata* and other *Caenorhabditis* genomes using either a *C. inopinata*-specific TE library (Kawahara *et al*., 2023) or a 14-species *Caenorhabditis* TE library (Sun *et al*., 2022). Assemblies were obtained from WormBase Parasite release 19 for *C. elegans* (PRJNA13758), *C. nigoni* (PRJNA384657), *C. sinica* (PRJNA194557), *C. brenneri* (PRJNA20035), *C. remanei* (PRJNA577507), *C. quiockensis* (PRJEB11354), *C. latens* (PRJNA248912), *C. angaria* (PRJNA51225), *C. bovis* (PRJEB34497), *C. briggsae* (PRJNA784955), *C. niphades* (PRJEB53466), and *C. drosophilae* (PRJEB67483). The *C. japonica* v2 assembly was generated in our laboratory.

Self-alignments were generated with LAST v1542-1 (Camacho *et al*., 2009) using lastdb - uNEAR and lastal. MAF (Multiple Alignment Format) output was parsed and visualised with ggplot2 v4.0.0 (https://ggplot2.tidyverse.org/) implemented in R v4.4.1 (https://www.r-project.org/). Forward–forward matches were plotted along the main diagonal, whereas forward–reverse matches were plotted along the anti-diagonal.

### Genome screening for *PIF/Harbinger* elements

*PIF/Harbinger* elements were identified in the *C. inopinata* genome with seqkit locate (Shen *et al*., 2016) by searching for lineage-specific 15-bp TIR sequences, allowing up to two mismatches (-m 2). Candidate elements were defined as oppositely oriented TIR-like pairs located within 4 kb of each other. Candidates lacking identical 3-bp flanking sequences at both ends were excluded. When nested or overlapping pairs were detected within the same interval, the shortest candidate was retained. Terminal sequences of curated elements were aligned with MAFFT v7.505 (Katoh and Standley, 2013) (--auto --adjustdirection) and used to summarise TIR motifs with WebLogo v3.7.9 (Crooks *et al*., 2004). Terminal windows comprised 80 nt from each end for *Turmoil2*-like elements and 30 nt from each end for *Turmoil1*-like elements. TSD motifs were summarised from the 3-bp flanking sequences.

ORFs within identified *PIF/Harbinger* elements were predicted with EMBOSS getorf (v6.6.0.0) (Rice *et al*., 2000). Predicted proteins were scanned with pfam_scan.pl (v1.6.4) (Mistry *et al*., 2021) for DDE transposase domains (DDE_Tnp_4; PF13359.11) and MADF DNA-binding domains (MADF_DNA_bdg; PF10545.14). Elements were classified as MADF-only when a MADF_DNA_bdg-containing ORF was detected in the absence of a DDE_Tnp_4-containing ORF, as DDE-only when the reciprocal pattern was observed, and as non-autonomous MITEs when neither domain was detected.

### Clustering analysis of *PIF/Harbinger* elements

Exact duplicate sequences were removed with vsearch v2.18.0 (Rognes *et al*., 2016) using --derep_fulllength --sizeout --minuniquesize 1, and dereplicated sequences were clustered at 80% identity with --cluster_size --id 0.80 --strand both --sizein. Clusters containing at least 10 sequences were retained. Sequences within each cluster were aligned with MAFFT v7.505 (Katoh and Standley, 2013) (--auto --adjustdirection) and trimmed with trimAl v1.4 (Capella-Gutierrez *et al*., 2009) (-gt 0.8). Pairwise nucleotide divergence was calculated under the Kimura two-parameter model with EMBOSS distmat (v6.6.0.0) (Rice *et al*., 2000) (-nucmethod 2). Summary statistics were calculated from unique pairwise distances in the upper triangle of each distance matrix. The weighted median element length for each cluster was calculated from dereplication copy counts. Sequence similarity to Ci_Harb_255 was assessed by BLASTN with BLAST+ v2.12.0 (Camacho *et al*., 2009) using -dust no -evalue 1e-5 -word_size 10 -outfmt 6.

### Phylogenetic analysis

Amino-acid sequences were aligned with MAFFT v7.505 (--auto) (Katoh and Standley, 2013) and trimmed with trimAl v1.4 (-automated1) (Capella-Gutierrez *et al*., 2009). Maximum-likelihood phylogenies were inferred with IQ-TREE v2.1.4 (Minh *et al*., 2020), with substitution models selected by ModelFinder (Kalyaanamoorthy *et al*., 2017) and branch support assessed using 1,000 SH-aLRT replicates and 1,000 ultrafast bootstrap replicates. Sequences used in the analysis are listed in Supplementary Table 1. For conserved-domain comparisons within Caenorhabditis, Pfam-annotated domain regions were extracted from the protein sequences, aligned with MAFFT (--auto), and visualised with ggmsa v1.10.0 (Zhou *et al*., 2022) in R v4.4.1.

### RNA-seq analysis

RNA-seq data from six libraries (two biological replicates each for egg, male, and female samples) were retrieved from GenBank (BioProject PRJDB5687). Reads were aligned to the *C. inopinata* reference genome using STAR (v2.7.11b) (Dobin *et al*., 2013) with a curated gene annotation (v7.11; Supplementary File 1). To retain multi-mapping reads, particularly those derived from repetitive TE-related sequences, mapping thresholds were relaxed (--outFilterMultimapNmax 200 and --winAnchorMultimapNmax 200). Read counts were quantified using TEcount (TEtranscripts v2.2.3) (Jin *et al*., 2015), which incorporates multi-mapping reads during expression estimation. Although TE annotations generated from RepeatMasker output using a *C. inopinata*-specific TE library were used during quantification, only gene-level count data were used in the present analyses. Raw gene count tables were imported into R, where library sizes were normalised using calcNormFactors (edgeR v4.2.2), and expression levels were converted to counts per million (CPM).

### MADF domain diversity

MADF-domain regions were extracted from protein sequences, aligned with MAFFT v7.505 (-- localpair --maxiterate 1000), and trimmed with trimAl (v1.4; -automated1) to obtain domain-only alignments. Pairwise protein distances under the Kimura model were computed using PHYLIP protdist (PHYLIPNEW v3.69.650; EMBOSS v6.5.7.0) (Alzohairy, 2009). To avoid overweighting identical copies, zero-distance clusters within each family were collapsed by single-linkage hierarchical clustering (cut height *h* ≤ 1×10⁻¹²), retaining one representative per cluster (yielding n = 7 for each family). We then performed bootstrap resampling with replacement (10,000 replicates; each replicate sampled *k* = 7 sequences). For each replicate, the mean of the upper triangle of the symmetrised Kimura distance matrix was calculated; uncertainty was summerised as 95% percentile bootstrap confidence intervals.

## Results

### Isolation of a spontaneous Dpy mutant in *C. inopinata*

A spontaneous Dpy mutant of *C. inopinata* with reduced body size was isolated from a laboratory culture of the wild-type strain NKZ35 (Fig. 1A). Genetic crossing demonstrated that this phenotype was caused by a single recessive mutant allele, designated *tj217*. Whereas homozygous *tj217* animals derived from heterozygous parents displayed a Dpy phenotype, homozygous animals derived from homozygous mothers exhibited a more severe Dpy phenotype (extreme Dpy, or Ex-Dpy) (Fig. 1A). These results indicate that maternal contribution of the relevant gene influences the severity of the Dpy phenotype.

**Fig. 1.**
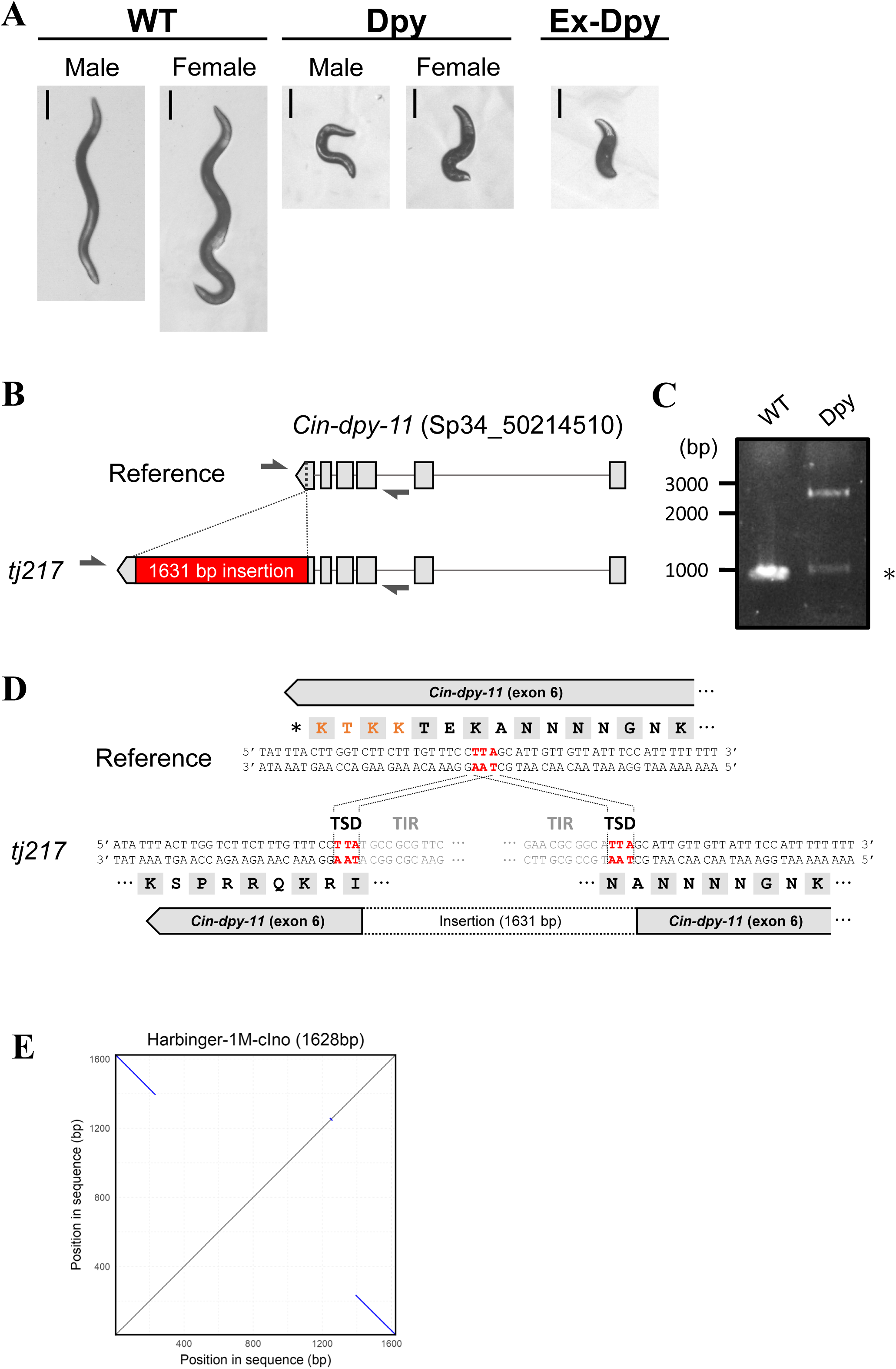
Identification of a *PIF/Harbinger-*family insertion in the coding region of *Cin-dpy-11.* (A) Isolation of a spontaneous dumpy (Dpy) mutant in *C. inopinata*. Adult worms of wild-type (WT), Dpy (*tj217* homozygotes derived from heterozygous mothers), Ex-Dpy (extreme-Dpy; a *tj217* homozygote derived from a *tj217* homozygous mother) are shown. Scale bars, 200 μm. (B) Schematic structures of the WT and the *tj217* allele of *Cin-dpy-11*. A 1,631-bp insertion in exon 6 of the *tj217* allele is indicated (exons are shown as grey boxes). Arrows indicate the positions of the primers used in (C). (C) PCR analysis of the *Cin-dpy-11* (Sp34_50214510) gene regions in WT and Dpy. An asterisk indicates a PCR-artifact. (D) Sequences of the WT and *tj217* allele around the insertion site. Three-base target site duplications (TSDs) flank the insertion, and the terminal inverted repeats (TIRs) of the *PIF/Harbinger* element are present at both ends (only the terminal 10 bp of each TIR are shown). The C-terminal ER-retention motif (KKTK) of DPY-11 is shown in orange and is disrupted by the insertion. (E) Self dot-plot alignment of *Harbinger-1M_cIno* sequences.

### Insertion of *PIF/Harbinger* element in *Cin-dpy-11*

To identify the mutation underlying the *tj217* Dpy phenotype, we performed whole-genome sequencing on individual Dpy and wild-type worms. Sequencing of five single-worms from each genotype yielded genome coverage of 130–380× per sample. We then screened for variants that were consistently homozygous in Dpy animals and absent from wild-type controls. Among these variants, we identified an insertion in *Cin-dpy-11* (*Sp34_50214510*), a one-to-one orthologue of *C. elegans dpy-11* (Fig. 1B). Because loss of *dpy-11* causes a recessive dumpy phenotype in *C. elegans* (Ko and Chow, 2002; Vora et al., 2025), the insertion in *Cin-dpy-11* is a strong candidate for the causal lesion underlying *tj217*. *Cin-dpy-11* comprises six exons, spans approximately 3.2 kb of genomic sequence and shares 93.9% amino acid identity (231/246 residues) with *C. elegans* DPY-11 (Supplementary Fig. 1).

We validated the candidate insertion by PCR using genomic DNA from wild-type and Dpy worms. The wild-type template yielded a single ∼1-kb amplicon, whereas the Dpy template yielded a larger ∼2.6-kb product (Fig. 1C). Sequencing of the mutant amplicon revealed a 1,631-bp insertion within exon 6 of *Cin-dpy-11*. The insertion disrupts the C-terminal KKTK motif of DPY-11, which is predicted to contribute to ER retention, thereby providing a plausible molecular basis for loss of function (Fig. 1D; Supplementary Fig. 1). Sequence analysis further showed that the inserted sequence corresponds to a *PIF/Harbinger*-family DNA transposon with TIRs and a flanking 3-bp TSD (TTA) (Fig. 1D and 1E). We designated this element *Harbinger-1M_cIno*. Members of the *PIF/Harbinger* superfamily typically encode two proteins required for transposition: a DDE-family transposase and a protein with a MADF DNA-binding domain. Annotation of the predicted ORFs within *Harbinger-1M_cIno* identified a MADF DNA-binding domain but did not detect a recognisable DDE-transposase domain, suggesting that *Harbinger-1M_cIno* is non-autonomous.

### Genome-wide distribution of the *Harbinger-1M_cIno* family

Having identified *Harbinger-1M_cIno* as the element inserted in *Cin-dpy-11*, we next asked whether it belongs to a broader element family in the *C. inopinata* genome. To address this, we searched the genome using the terminal 15-bp TIR motif of *Harbinger-1M_cIno* together with filters for inverted terminal matches, TSDs, and element length. This search recovered 258 related elements, which we hereafter refer to as the *Harbinger-1M_cIno* family (Supplementary Table 2). For convenience, the recovered elements were indexed by size as *Ci_Harb_1* to *Ci_Harb_258*, with *Ci_Harb_258* denoting the longest element. These elements ranged from 180 to 2,752 bp in length, with a median of 394 bp, and most (191/258) were shorter than 500 bp (Fig. 2A). A distinct peak was present at approximately 1.63 kb, corresponding to the size of *Harbinger-1M_cIno* itself. Conserved terminal blocks were evident at both ends, and the elements were flanked by characteristic 3-bp TSDs with a TWA consensus, most commonly TTA or TAA (Fig. 2B and 2C). The family was distributed predominantly on the autosomes, with only three copies detected on the X chromosome (Fig. 2D). This bias resembles the genomic distribution previously reported for the *C. inopinata* DNA transposons mCi-hAT1 and mCi-hAT4 (Hatanaka *et al*., 2024). Together, these results show that the element inserted in *Cin-dpy-11* is one member of a genome-wide *PIF/Harbinger*-derived family in *C. inopinata*.

**Fig. 2.**
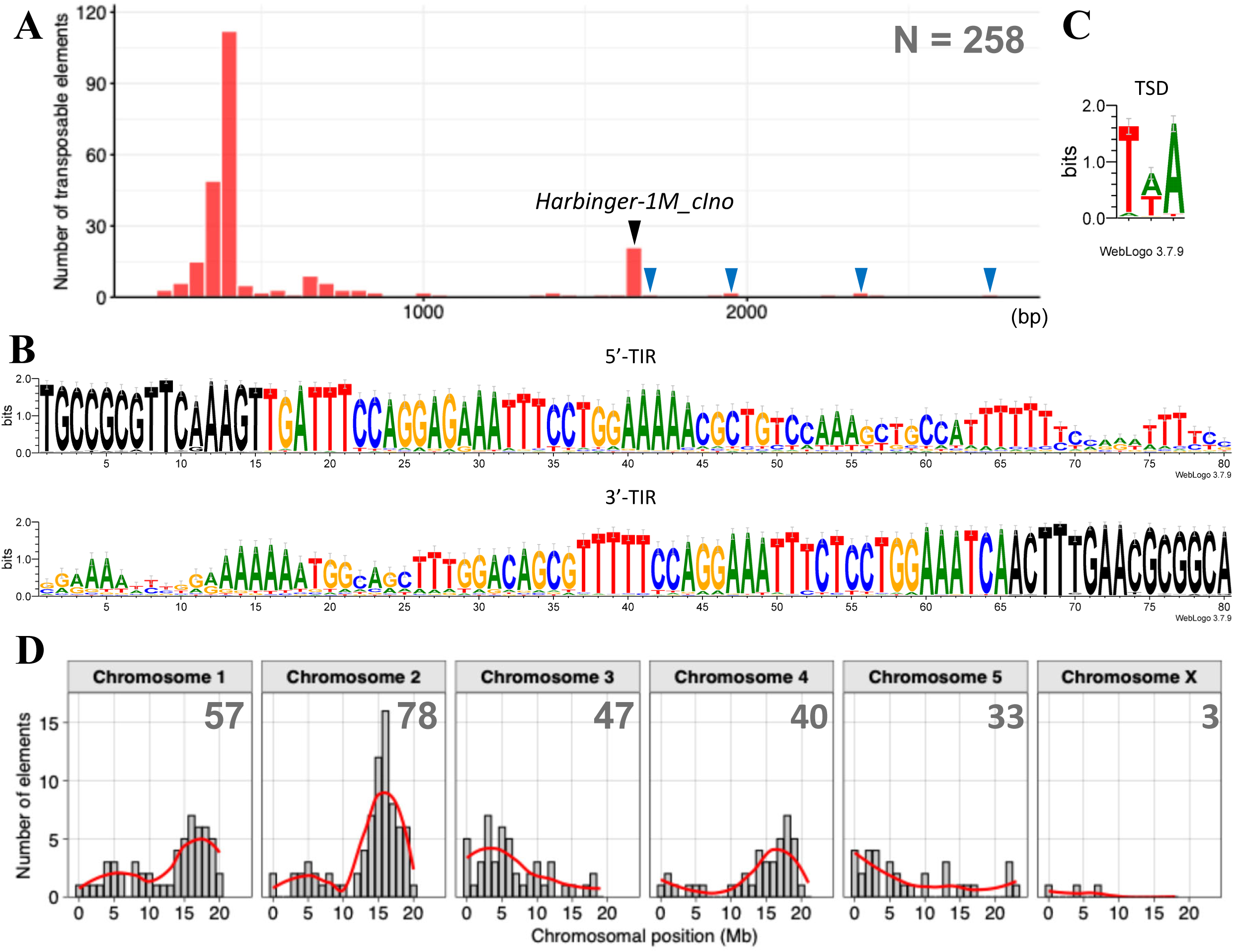
Characterisation of *Harbinger-1M_cIno*–like transposable elements in the *C. inopinata* genome. (A) Histogram of lengths for 258 *Harbinger-1M_cIno*–like transposable elements (TEs) extracted from the *C. inopinata* genome. The x-axis shows TE length, and the y-axis shows number of TEs per size bin. Elements with a DDE transposase domain are marked by blue arrowheads. Those near 1.63 kb, corresponding to *Harbinger-1M_cIno* is indicated by black arrowheads. (B) Sequence logos for 80-bp TIRs at both ends. Black traces indicate query sequences used for TE extraction. (C) Sequence logos for 3-bp TSDs. (D) Chromosomal distribution of *Harbinger-1M_cIno*-like TEs. TE counts per chromosome are shown at the top right.

A notable feature of this size distribution was that coding capacity was concentrated in the relatively small subset of long elements, particularly those around 1.6 kb, whereas the much more abundant short elements were predominantly noncoding derivatives. To examine this explicitly, we annotated predicted ORFs across all 258 family members. Pfam searches identified five elements carrying only DDE transposase domains (*Ci_Harb_250*, *Ci_Harb_252*, *Ci_Harb_253*, *Ci_Harb_255*, and *Ci_Harb_256*), 19 elements carrying only MADF domains, and one element, *Ci_Harb_258*, carrying both DDE and MADF domains (Fig. 3A; Supplementary Tables 2 and 3). Thus, long-elements clustered around 1.63 kb and above retained recognisable protein-coding potential, whereas most shorter elements appear to represent non-autonomous or degenerate derivatives. After manual curation, however, we found that only one DDE-bearing element, *Ci_Harb_252*, retained an apparently intact transposase gene, whereas the remaining DDE-like loci were pseudogenised. By contrast, 14 of the 19 MADF-bearing loci retained intact MADF domains. *Ci_Harb_258* was particularly informative because, as the longest element and the only family member carrying both DDE- and MADF-related loci, it provides the clearest structural reference for the family; however, both loci in *Ci_Harb_258* were pseudogenised. In particular, the MADF-related locus carries a 10-bp deletion in exon 2 that causes a frameshift and disrupts most of the MADF domain (Fig. 3A and 3B; Supplementary Fig. 2). RNA-seq analysis further showed that the intact DDE transposase gene *Sp34_30019800* and several MADF-domain genes were expressed across eggs, adult males, and adult females, whereas most pseudogenes showed little or no expression (Fig. 3C and 3D). These observations indicate that the *Harbinger-1M_cIno* family contains only a small number of potentially functional coding loci embedded within a much larger population of short, noncoding derivatives.

**Fig. 3.**
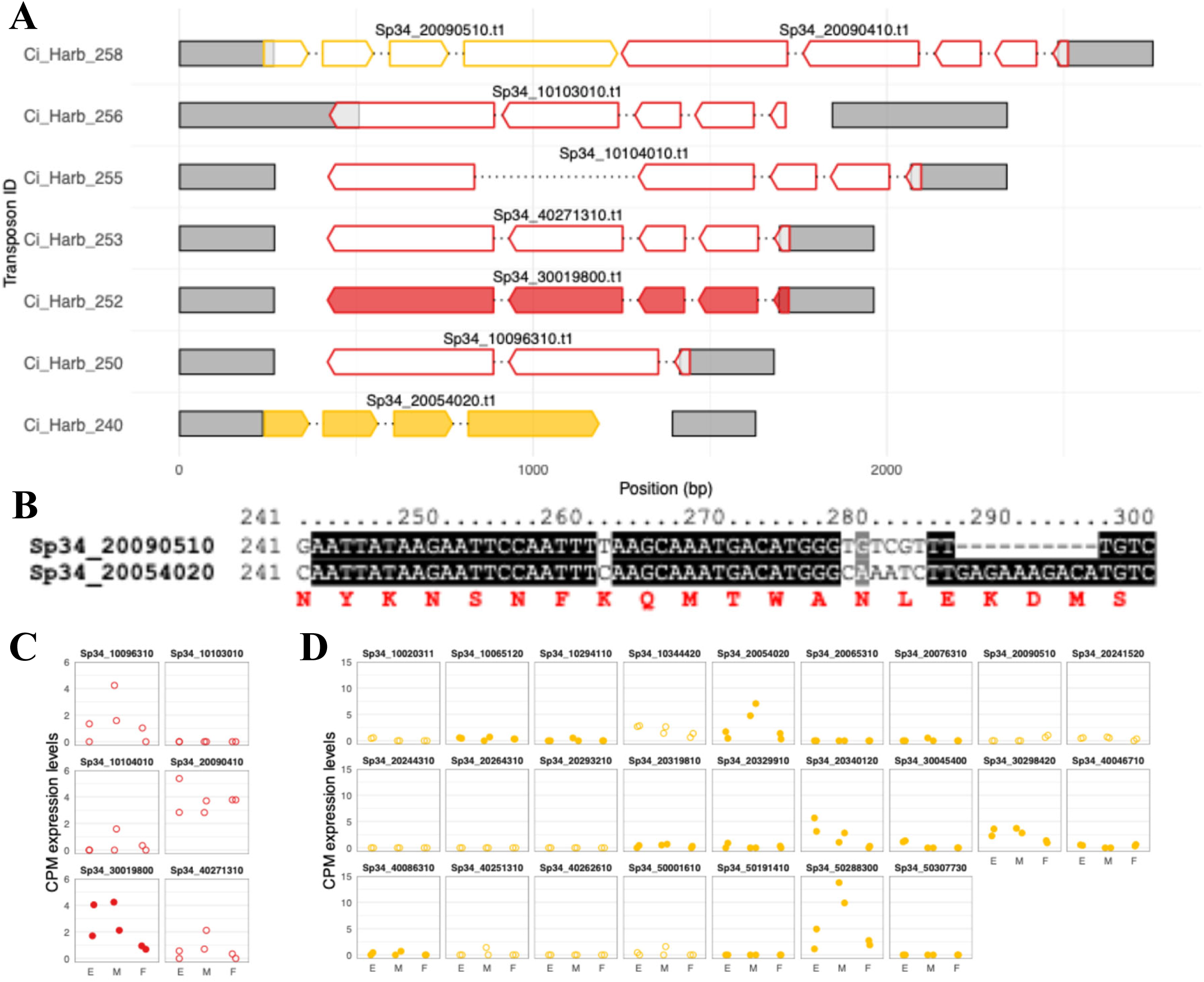
Structural organisation and expression of DDE- and MADF-domain loci within the *Harbinger-1M_cIno* family. (A) Schematic structures of *Harbinger-1M_cIno* family members carrying DDE transposase- or MADF-domain loci. Red and gold gene models indicate DDE- and MADF-domain loci, respectively; filled models denote predicted intact genes and open models denote pseudogenes. Grey boxes indicate putative TIRs. (B) Alignment of the MADF-coding region in *Ci_Harb_240* (*Sp34_20054020*) and *Ci_Harb_258* (*Sp34_20090510*). A 10-bp deletion in exon 2 of *Sp34_20090510* causes a frameshift predicted to disrupt most of the MADF domain. (C) Expression of DDE-bearing loci across eggs (E), adult males (M), and adult females (F). (D) Expression of MADF-bearing loci across the same stages. In (C) and (D), each facet represents a single locus; filled circles indicate predicted genes and open circles indicate pseudogenes. All panels share the same y-axis scale (counts per million, CPM).

We next examined how these coding and noncoding derivatives are related to one another. Sequence-similarity clustering of the 258 elements resolved 74 groups, indicating substantial diversification within the family. Because most groups were small, we focused on the four largest clusters for further analysis. One major cluster comprised elements closely similar to *Harbinger-1M_cIno* in both size and sequence and showed relatively low divergence, whereas the other large clusters consisted of shorter and more heterogeneous elements with substantially greater divergence (Fig. 4A). We then used *Ci_Harb_258* as a structural reference to compare similarity across the family. Most elements retained highly similar terminal regions, but similarity across the internal region was partitioned. One group preferentially retained the segment corresponding to the MADF-associated region of *Ci_Harb_258*, whereas the DDE-bearing elements retained the segment corresponding to the transposase-encoding portion (Fig. 4B). Together, these patterns are consistent with diversification from a common *Harbinger-1M_cIno*–like ancestor through internal deletion and related structural rearrangements.

**Fig. 4.**
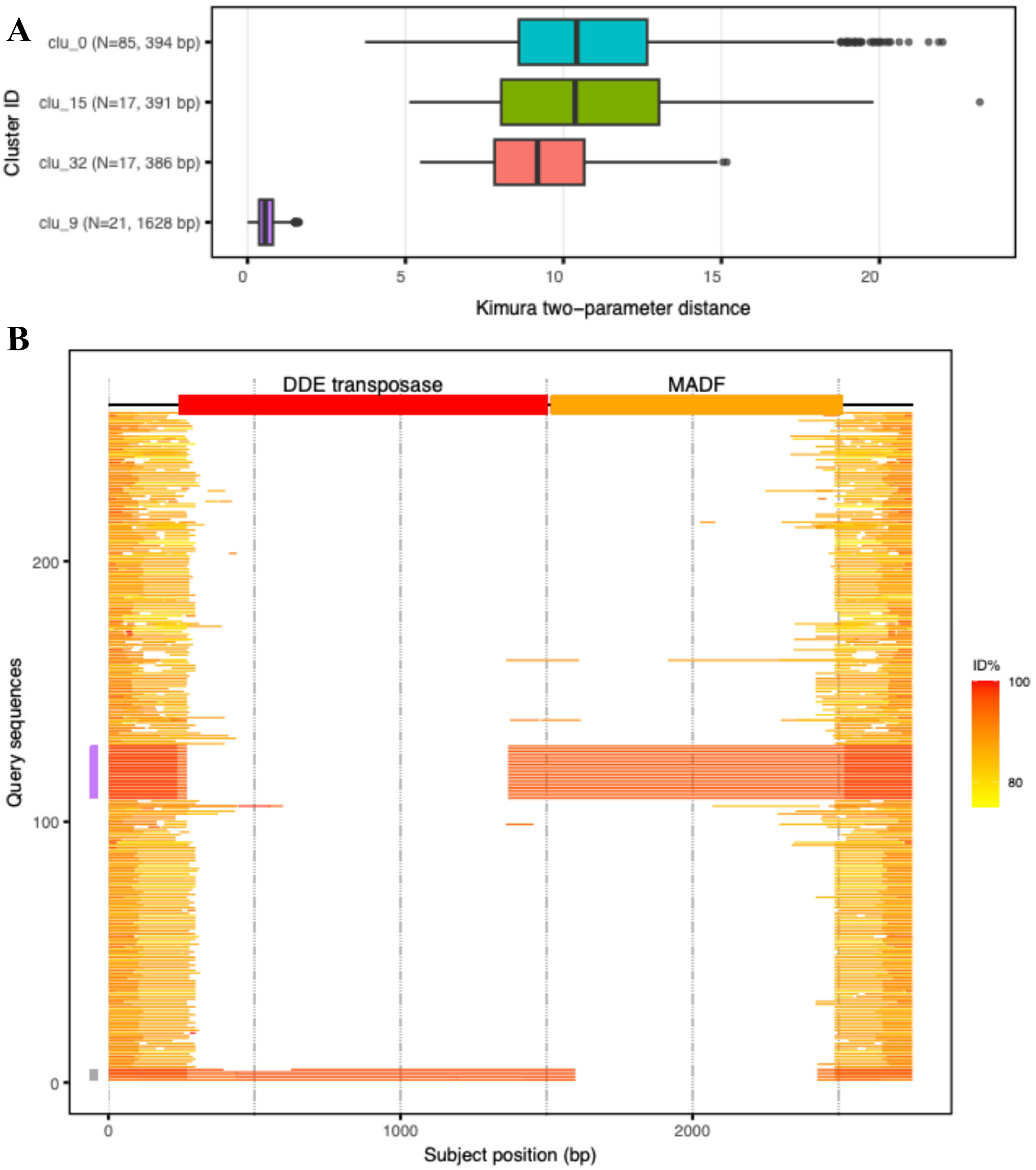
Divergence and structural partitioning within the *Harbinger-1M_cIno* family. (A) Kimura two-parameter distances within the four largest sequence clusters. Cluster labels indicate the number of elements and median length. The long-element cluster (clu_9; median 1,628 bp) shows markedly lower divergence than the shorter clusters. (B) BLAST similarity of *Harbinger-1M_cIno* family members to *Ci_Harb_258*. Each horizontal line represents one query sequence; colors indicate nucleotide identity. Most elements retain high similarity at the termini, whereas internal similarity is partitioned between the DDE-transposase and MADF-associated regions. Gray and purple side bars indicate DDE-bearing elements and clu_9 members, respectively.

### Turmoil2 affiliation of the Harbinger-1M_cIno family

Because the 258-element search was anchored on the TIR sequence of *Harbinger-1M_cIno*, it was designed to recover the family related to the inserted element rather than to comprehensively identify all *PIF/Harbinger* lineages in *C. inopinata*. We therefore next asked how the *Harbinger-1M_cIno* family relates to the *PIF/Harbinger* elements in *C. elegans* (named *Turmoil1* and *Turmoil2*). DDE transposase phylogeny across *Caenorhabditis* and related *PIF/Harbinger* elements resolved two major clades corresponding to *Turmoil1* and *Turmoil2* and recovered the conserved D-D-E catalytic motif in both groups (Fig. 5). Within this framework, the coding component of the *Harbinger-1M_cIno* family is associated with the *Turmoil2* lineage.

**Fig. 5.**
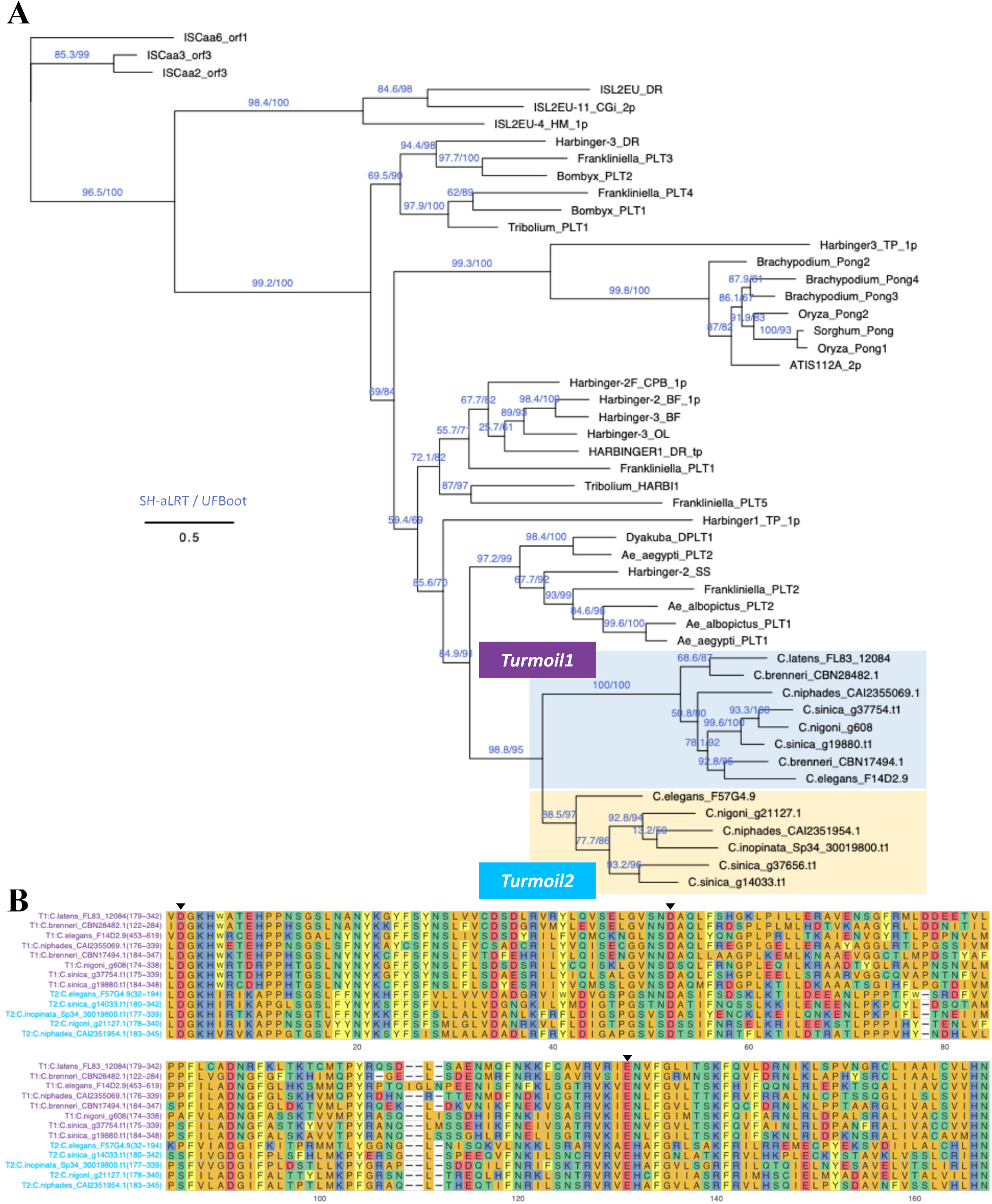
Phylogenetic relationships of *PIF/Harbinger* superfamily transposases. (A) Maximum-likelihood phylogeny inferred from DDE transposase sequences of *Caenorhabditis* species together with representative *PIF/Harbinger* elements from other taxa. Numbers above branches indicate SH-aLRT and ultrafast bootstrap (UFBoot) support values. Transposases from the ISL2 and ISL2EU groups were used as outgroups. (B) Multiple sequence alignment of the DDE domain in *Turmoil1*- and *Turmoil2*-derived transposases. Numbers in parentheses indicate amino acid positions in the full-length proteins. Conserved DDE catalytic residues are marked by arrowheads.

For comparative purposes, we then surveyed 14 *Caenorhabditis* genomes using a common repeat library containing *Turmoil1*- and *Turmoil2*-type consensuses. Copy number and family composition varied markedly among species (Supplementary Fig. 3; Supplementary Table 4). *C. inopinata*, *C. bovis*, and *C. japonica* each contained at least 80 detectable *PIF/Harbinger* hits, whereas the remaining 11 species contained 15 or fewer. Within this comparative survey, most detectable hits in *C. inopinata* matched the *Turmoil2*-type consensus, whereas *C. bovis* and *C. japonica* were dominated by *Turmoil1*-type hits. Outside *C. inopinata*, *PIF/Harbinger* insertions typically retained both DDE- and MADF-related genes within the same TIR-bounded locus or were represented by DDE-only loci; by contrast, MADF-only loci were not detected. Representative examples from *C. niphades* and *C. nigoni* showed the canonical head-to-head arrangement of DDE and MADF genes, closely resembling the structure of *Ci_Harb_258* (Supplementary Fig. 3E). Together, these results indicate that the extensive proliferation of MADF-only derivatives associated with the *Harbinger-1M_cIno* family is unusual among the *Caenorhabditis* genomes examined here and is especially pronounced in *C. inopinata*.

### A Turmoil1-derived PIF/Harbinger family in C. inopinata

The *Turmoil*-based comparison suggested that *C. inopinata* might retain a second *PIF/Harbinger*-derived family distinct from *Harbinger-1M_cIno*. To examine this directly, we analysed a species-specific repeat library for *C. inopinata*, which identified three *PIF/Harbinger*-derived consensus families. One showed clear similarity to *C. elegans Turmoil1*, whereas the other two were derived from the *Harbinger-1M_cIno* family (Supplementary File 2). TBLASTN searches using *C. niphades Turmoil1* transposase (CAI2355069.1) and MADF (CAI2355070.1) as queries recovered seven MADF-like loci but no DDE transposase-like loci, and all seven MADF loci retained full-length MADF domains. Thus, unlike the *Harbinger-1M_cIno* family, which includes a small subset of long elements with DDE and/or MADF coding potential, this second family was represented in the current genome only by MADF-bearing loci.

We next examined whether these MADF loci retained recognisable terminal structures. Inverted-repeat scans of the ±1 kb regions flanking each MADF gene detected matching flanking repeats at four of the seven loci, although these were shorter and less conserved than those associated with the *Harbinger-1M_cIno* family. These observations indicate that *C. inopinata* retains a distinct *Turmoil1*-like MADF family, whereas the corresponding DDE transposase has been lost and the flanking TIRs have become degraded. To define this family more explicitly, we reconstructed a consensus from the conserved terminal sequences shared by the seven MADF-coding loci and by additional noncoding loci that retained the same terminal architecture. We designated the resulting *Turmoil1*-derived family *Harbinger-2M_cIno*, to distinguish it from the *Turmoil2*-associated *Harbinger-1M_cIno* family.

### Contrasting histories of the two *C. inopinata PIF/Harbinger* families

We then compared the evolutionary histories of the two *PIF/Harbinger*-derived families in *C. inopinata*, *Harbinger-1M_cIno* and *Harbinger-2M_cIno*. Reconstruction of a *Turmoil1*-derived consensus followed by genome-wide searching identified 1,376 *Harbinger-2M_cIno* copies ranging from 80 to 2,878 bp in length, with a median size of 138 bp, distributed predominantly on autosomes (Fig. 6; Supplementary Table 5). Although these elements retained 3-bp TWA target-site duplications, their terminal and internal sequences were less conserved than those of *Harbinger-1M_cIno* (Fig. 6B and 6C). Divergence analyses likewise showed broader sequence variation and older substitution profiles, lacking the low-divergence peak characteristic of *Harbinger-1M_cIno* (Fig. 6E). Together, these features indicate that *Harbinger-2M_cIno* is older and more degenerate than *Harbinger-1M_cIno*.

**Fig. 6.**
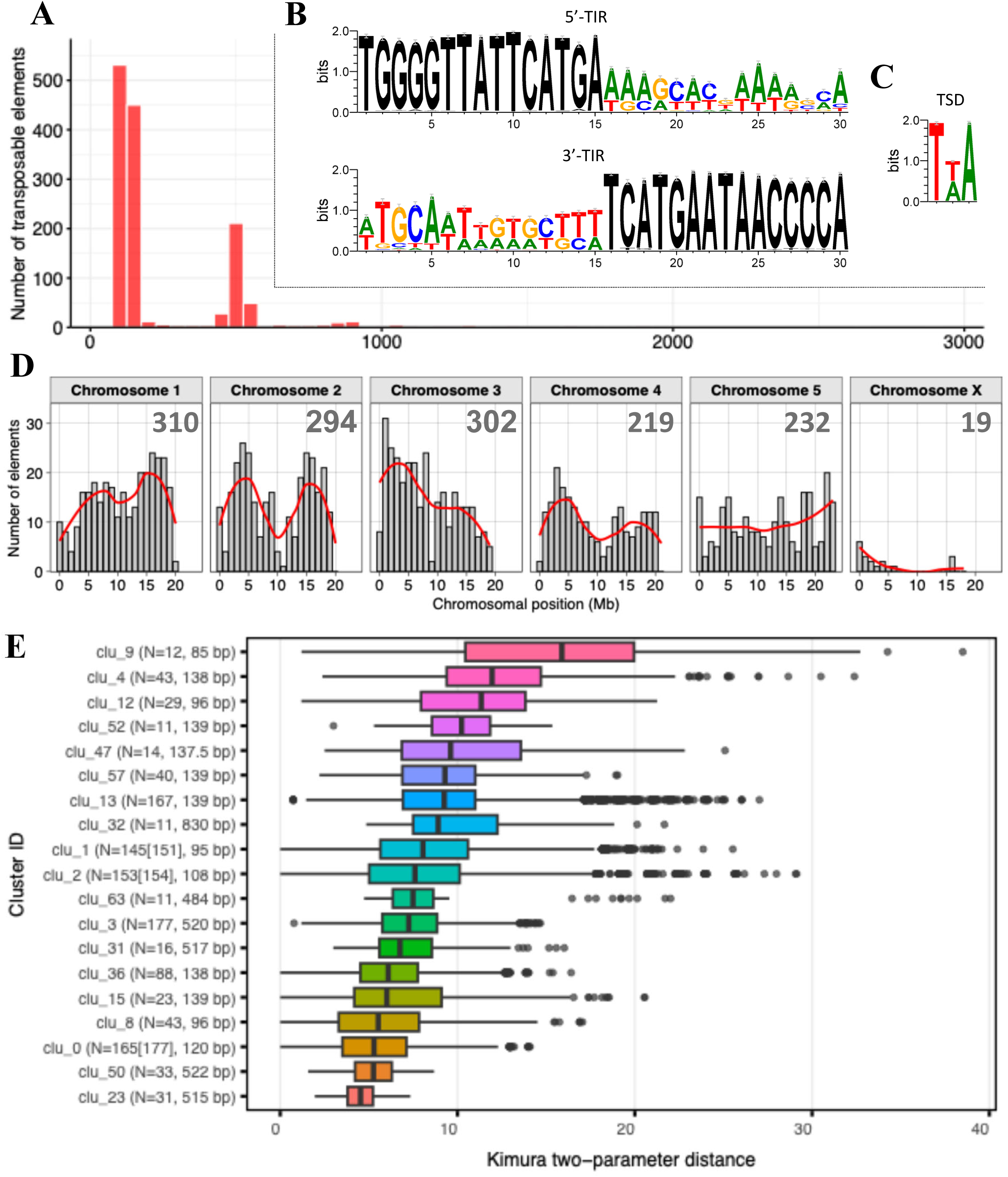
Genome-wide characterisation of the *Harbinger-2M_cIno* family in *C. inopinata*. (A) Length distribution of 1,376 *Harbinger-2M_cIno* copies. (B) Sequence logos of the 5′ and 3′ TIRs. (C) Sequence logo of the 3-bp TSD. (D) Chromosomal distribution of *Harbinger-2M_cIno* copies across the six chromosomes of *C. inopinata*. Numbers indicate total copies per chromosome. (E) Kimura two-parameter distances within the major sequence clusters of the family. Cluster labels indicate cluster ID, number of unique elements [total number of elements], and median length.

A direct comparison of *Turmoil1*- and *Turmoil2*-derived MADF genes supported the same conclusion. Phylogenetic analysis of 21 MADF loci, including seven *Turmoil1*-derived and 14 *Turmoil2*-derived sequences, resolved two reciprocally monophyletic clades, with markedly longer branches in the *Turmoil1*-derived clade (Supplementary Fig. 4A and 4B). Bootstrap estimates of mean pairwise Kimura protein distance were 0.143 (95% CI, 0.082–0.189) for *Turmoil1* and 0.026 (95% CI, 0.016–0.033) for *Turmoil2*, indicating substantially greater divergence in the *Turmoil1*-derived MADF set (Supplementary Fig. 4C). RNA-seq analysis further detected transcription of all seven *Turmoil1*-derived MADF genes, without a strong stage-specific bias (Supplementary Fig. 4D). Together, these results indicate that the two *PIF/Harbinger*-derived families in *C. inopinata* have distinct histories: *Harbinger-1M_cIno* is comparatively homogeneous and consistent with more recent proliferation, whereas *Harbinger-2M_cIno* is more deeply diverged and structurally degraded, although its MADF genes remain transcriptionally retained.

## Discussion

The main advance of this study is not simply the identification of a transposon insertion in *Cin-dpy-11*, but the recognition that this lesion reveals an unusual evolutionary configuration of a *PIF/Harbinger*-derived family in *C. inopinata*. Canonical autonomous *PIF/Harbinger* elements generally encode both DDE transposase and MADF functions within the same element (Kapitonov and Jurka, 1999; Sinzelle *et al*., 2008). Against that background, the *Turmoil2*-associated family *Harbinger-1M_cIno* is notable because these two functions are retained in separate descendant elements, yet only the MADF-bearing branch has undergone broad genome-wide expansion, whereas DDE-bearing descendants remain restricted to a small number of copies. We suggest that the key signal here is not modularity itself, which is a general feature of *PIF/Harbinger* biology, but a markedly asymmetric pattern of descendant-family expansion. In this sense, the *tj217* mutant reveals not only a causal lesion but also a naturally occurring example of how a canonical two-ORF *PIF/Harbinger* family can diversify into unequally expanded descendant branches.

The interpretation that the *Cin-dpy-11* insertion underlies the Dpy phenotype is biologically compelling. In *C. elegans*, *dpy-11* encodes a membrane-associated thioredoxin-like protein required for normal body and sensory-organ morphogenesis (Ko and Chow, 2002; Vora *et al*., 2025), making disruption of its coding region fully consistent with the strong body-shape defect observed here. Although formal proof by rescue or targeted reversion remains to be obtained, the convergence of genetic mapping, structural validation, and functional correspondence to *C. elegans dpy-11* strongly supports the insertion as the causal lesion.

The abundance of short, domainless derivatives in the *Harbinger-1M_cIno* family is also consistent with a MITE-like mode of family expansion. MITEs are short non-autonomous DNA transposons that commonly arise through internal deletion of larger autonomous elements while retaining terminal features required for mobilisation (Yang *et al*., 2009). In this context, the many short *Harbinger-1M_cIno* derivatives are most naturally interpreted as MITE-like noncoding descendants of a more complete ancestral element, and their predominance over the longer coding-capable copies suggests that amplification of such noncoding derivatives has been a major component of *Turmoil2*-associated family evolution in *C. inopinata*. Against this background, the coding derivatives represent only a minor subset of the family. Within that subset, MADF-bearing descendants appear to have persisted and expanded more successfully than DDE-bearing descendants, whereas the great majority of copies are short noncoding derivatives rather than coding descendants.

By contrast, the *Turmoil1*-derived family in *C. inopinata* appears to represent a later stage of erosion rather than a parallel active lineage. It is retained as MADF-bearing loci without a detectable DDE counterpart, its terminal structures are more degraded, and its sequences are more diverged. For that reason, we view the *Turmoil1*-derived family mainly as a useful contrast that defines the range of evolutionary states present in the same genome. Its retained transcription is intriguing, but transcription alone does not establish present-day function. At most, these loci may represent substrates for subsequent host co-option, as seen for *PIF/Harbinger*-derived proteins in other animal and plant lineages (Kapitonov and Jurka, 1999; Sinzelle *et al*., 2008; Velanis *et al*., 2020). More importantly, the *Turmoil1*-derived family no longer behaves like the *Turmoil2*-associated family and therefore is unlikely to contribute to the ongoing activity inferred for *Harbinger-1M_cIno*.

Comparative analysis also suggests that *Turmoil* elements have followed different structural trajectories in *C. elegans* and *C. inopinata*. Relative to the compact terminal structures typical of *PIF/Harbinger* elements, the *Turmoil1* and *Turmoil2* elements described in *C. elegans* appear to carry unusually extended, composite TIR structures, whereas the *Turmoil2*-associated family in *C. inopinata* retains shorter termini resembling those seen in other *Caenorhabditis* species. Together with the TE-rich genome of *C. inopinata* and its loss of several small-RNA pathway components, this contrast raises the possibility that lineage-specific host–TE interactions have shaped *Turmoil* architecture and activity differently in the two species. At present, however, this remains a hypothesis rather than a demonstrated mechanism.

The spontaneous insertion into *Cin-dpy-11* indicates that at least one *Turmoil2*-associated *PIF/Harbinger* element has been active under laboratory conditions in *C. inopinata*. Together with the TE-rich genome of this species (Kanzaki *et al*., 2018) and the recent activity of other DNA transposon families, including the Ci-hAT1/mCi-hAT1 system (Hatanaka *et al*., 2024), this observation suggests that *C. inopinata* provides a host background permissive for recent transposon activity. However, this permissiveness should not be interpreted as a simple loss of TE control. Although *C. inopinata* has lost multiple components of the ERGO-1/26G-siRNA pathway (Kanzaki *et al*., 2018), many core small-RNA pathway genes remain conserved, implying that complementary silencing systems are still present. We therefore infer that TE surveillance in *C. inopinata* has been reconfigured rather than abolished. Within this altered host background, the two *PIF/Harbinger*-derived families appear to occupy different temporal states: *Harbinger-1M_cIno* remains linked to recent activity, whereas *Harbinger-2M_cIno* preserves the genomic signature of substantial past amplification but now appears to have lost present-day mobilisation competence. In this framework, host-side permissiveness may help explain why recent transposition is still detectable in *C. inopinata*, whereas family-specific properties, such as compatibility between DDE-bearing and MADF-bearing derivatives and recognition of family-specific terminal sequences, are more likely to determine which descendant classes continue to expand.

Several limitations define the next steps. First, the causal role of the *Cin-dpy-11* insertion, although strongly supported by the convergence of phenotype, structure, and gene identity, remains to be verified by rescue or targeted reversion. Second, the inference that DDE and MADF functions may be supplied in trans within the *Harbinger-1M_cIno* family remains hypothetical rather than mechanistically demonstrated. Third, the evolutionary placement of the shortest noncoding derivatives necessarily relies on indirect evidence from terminal structure and sequence similarity. These limitations are experimentally tractable. Direct mobilisation assays, DDE–MADF cross-complementation tests, and terminal-sequence recognition assays should determine whether the expanded MADF-bearing derivatives are still functionally linked to the residual DDE source.

In summary, our results support a model in which two *PIF/Harbinger*-derived families with a shared origin have followed sharply different trajectories in *C. inopinata*. The *Turmoil2*-associated family remains linked to recent activity, as evidenced by the *Harbinger-1M_cIno* insertion into *Cin-dpy-11*, whereas the *Turmoil1*-derived family appears more eroded, having lost a detectable DDE component and retained only degraded terminal structures. A distinguishing feature of *C. inopinata* is the broad proliferation of MADF-bearing derivatives, particularly in the *Turmoil2*-associated family, whereas a comparable genome-wide expansion of DDE-only derivatives is not evident. Together with the TE-rich genome of *C. inopinata*, the loss of multiple small-RNA pathway components, and recent activity of other DNA transposon families in this species, these findings are most consistent with a host background permissive for transposon activity. Within such a background, family-specific properties, including compatibility between DDE- and MADF-bearing derivatives and recognition of terminal sequences, are likely to influence which descendant classes remain mobilisation-competent and therefore expand. This framework makes *C. inopinata* a useful system for testing how host genome defense and *PIF/Harbinger* family architecture interact to shape lineage-specific transposon trajectories.

## Acknowledgements

Genome data analyses were performed in part using the DDBJ supercomputer system. We thank members of the Kikuchi laboratory for helpful discussions.

## Authors’ contributions

A.S. and T.K. conceptualised the study. S.O., K.K. and N.H. isolated and photographed a spontaneous mutant and performed genetic crosses. X.J. prepared a whole-genome sequencing library, and A.Y. performed whole-genome sequencing. S.O., K.K. and N.H. carried out PCR and validated the TE insertion by Sanger sequencing. X.J., K.S., and T.K. primarily conducted the computational analyses with support from S.S. X.J. and K.S. drafted the initial version of the manuscript, which was reviewed and edited by all authors. T.K. finalised the manuscript.

## Data Availability

The raw whole-genome sequencing reads generated in this study for Caenorhabditis inopinata strains SA1635 and SA1689 (five biological replicates each) have been deposited in the NCBI Sequence Read Archive (SRA) under BioProject accession number [PRJDB14908]. The wild-type (NKZ35, SA1689) and mutant (SA1635 [tj217]) nematode strains utilized in this work are available upon request from the corresponding author.

## Figure legends

**Supplementary Fig. 1. Conservation of the *dpy-11* gene in *C. elegans* and *C. inopinata*.**

Amino-acid alignment of DPY-11 proteins from *C. elegans* (F46E10.9) and *C. inopinata* (Sp34_50214510.t1), highlighting sequence conservation and the ER-retention signal (KKTK).

**Supplementary Fig. 2. Pseudogenisation of a MADF-domain gene by frameshift.**

(A) Pfam annotation identifies an intact MADF domain in Sp34_20054020.t1 (red), whereas the domain is truncated in Sp34_20090510.t1. (B) Nucleotide alignment of the MADF-coding regions of Sp34_20054020 and Sp34_20090510, showing a 10-bp deletion and the resulting disrupted reading frame. Exons of Sp34_20054020 are highlighted in green. The amino-acid translation of Sp34_20054020 is shown beneath the alignment.

**Supplementary Fig. 3. Detection of *Turmoil1*- and *Turmoil2*-type transposable elements in Caenorhabditis.**

(A) Copy numbers of *PIF/Harbinger*-like elements in 14 species based on cross-species RepeatMasker library. The families rnd-6_family-38341 and rnd-6_family-25689 are classified as *Turmoil1*-type, whereas rnd-6_family-8036 is *Turmoil2*-type. (B–D) Numbers of elements with DDE or MADF domains within 2 kb upstream or downstream of TE hits. Genome assemblies not at chromosome-scale are marked with asterisks. (E) Representative *Turmoil2*-like loci in *C. niphades* and *C. nigoni*, showing opposite-orientation DDE (red) and MADF (gold) genes flanked by grey TIRs.

**Supplementary Fig. 4. Greater divergence of MADF proteins in *Turmoil1* than in *Turmoil2*.**

(A) Maximum-likelihood phylogeny of MADF domains from *Caenorhabditis* species and representative *PIF/Harbinger* elements. Numbers above branches indicate SH-aLRT/UFBoot support for key nodes. Domesticated *PIF/Harbinger*-derived Myb/SANT proteins—NAIF1 (*Homo sapiens*) and ALP2 (*Arabidopsis thaliana*)—and a *Pangu* transposon from *Magallana gigas* were used as outgroups. (B) Multiple sequence alignment of MADF domains from *C. inopinata Turmoil1* (T1; purple, n = 7) and *Turmoil2* (T2; light blue, n = 14). Numbers in parentheses indicate amino acid positions in the full-length proteins. (C) Mean pairwise Kimura protein distances of MADF domains from *Turmoil1* and *Turmoil2*. Violin plots show bootstrap distributions of mean distances (10,000 replicates; n = 7 sequences per family after deduplication). Black points indicate bootstrap means; error bars indicate 95% percentile confidence intervals. A bootstrap test indicated greater divergence in *Turmoil1* than in *Turmoil2* (*p* < 0.001). (D) Expression of *Turmoil1*-derived MADF-domain genes across developmental stages (E, egg; M, male; F, female). Each facet represents one locus; points indicate two replicates per stage. All facets share the same y-axis scale (CPM).

## References

Alzohairy, A., 2009 Phylip and Phylogenetics. Genes, genomes and genomics.

Billi, A. C., S. E. J. Fischer and J. K. Kim, 2014 Endogenous RNAi pathways in C. elegans. WormBook: The Online Review of C. elegans Biology: 1–49.

Bourque, G., K. H. Burns, M. Gehring, V. Gorbunova, A. Seluanov et al., 2018 Ten things you should know about transposable elements. Genome Biology 19: 199–z.

Buckley, B. A., K. B. Burkhart, S. G. Gu, G. Spracklin, A. Kershner et al., 2012 A nuclear Argonaute promotes multigenerational epigenetic inheritance and germline immortality. Nature 489: 447–451.

Camacho, C., G. Coulouris, V. Avagyan, N. Ma, J. Papadopoulos et al., 2009 BLAST+: architecture and applications. BMC Bioinformatics 10: 421–421.

Capella-Gutierrez, S., J. M. Silla-Martinez and T. Gabaldon, 2009 trimAl: a tool for automated alignment trimming in large-scale phylogenetic analyses. Bioinformatics (Oxford, England) 25: 1972–1973.

Carver, T., S. R. Harris, M. Berriman, J. Parkhill and J. A. McQuillan, 2012 Artemis: an integrated platform for visualization and analysis of high-throughput sequence-based experimental data. Bioinformatics (Oxford, England) 28: 464–469.

Chen, X., O. Schulz-Trieglaff, R. Shaw, B. Barnes, F. Schlesinger et al., 2016 Manta: rapid detection of structural variants and indels for germline and cancer sequencing applications. Bioinformatics (Oxford, England) 32: 1220–1222.

Crooks, G. E., G. Hon, J. Chandonia and S. E. Brenner, 2004 WebLogo: a sequence logo generator. Genome Research 14: 1188–1190.

Danecek, P., J. K. Bonfield, J. Liddle, J. Marshall, V. Ohan et al., 2021 Twelve years of SAMtools and BCFtools. GigaScience 10: giab008.

Das, P. P., M. P. Bagijn, L. D. Goldstein, J. R. Woolford, N. J. Lehrbach et al., 2008 Piwi and piRNAs act upstream of an endogenous siRNA pathway to suppress Tc3 transposon mobility in the Caenorhabditis elegans germline. Molecular Cell 31: 79–90.

Dobin, A., C. A. Davis, F. Schlesinger, J. Drenkow, C. Zaleski et al., 2013 STAR: ultrafast universal RNA-seq aligner. Bioinformatics 29: 15–21.

Fattash, I., R. Rooke, A. Wong, C. Hui, T. Luu et al., 2013 Miniature inverted-repeat transposable elements: discovery, distribution, and activity. Genome 56: 475–486.

Guang, S., A. F. Bochner, D. M. Pavelec, K. B. Burkhart, S. Harding et al., 2008 An Argonaute transports siRNAs from the cytoplasm to the nucleus. Science (New York, N.Y.) 321: 537–541.

Han, M., C. Xiong, H. Zhang, M. Zhang, H. Zhang et al., 2015 The diversification of PHIS transposon superfamily in eukaryotes. Mobile DNA 6: 12–17. eCollection 2015.

Hancock, C. N., F. Zhang and S. R. Wessler, 2010 Transposition of the Tourist-MITE mPing in yeast: an assay that retains key features of catalysis by the class 2 PIF/Harbinger superfamily. Mobile DNA 1: 5–5.

Hatanaka, R., K. Tamagawa, N. Haruta and A. Sugimoto, 2024 The impact of differential transposition activities of autonomous and nonautonomous hAT transposable elements on genome architecture and gene expression in Caenorhabditis inopinata. Genetics 227.

Iwasaki, Y. W., K. Shoji, S. Nakagwa, T. Miyoshi and Y. Tomari, 2025 Transposon-host arms race: a saga of genome evolution. Trends Genet 41: 369–389.

Jin, Y., O. H. Tam, E. Paniagua and M. Hammell, 2015 TEtranscripts: a package for including transposable elements in differential expression analysis of RNA-seq datasets. Bioinformatics 31: 3593–3599.

Jurka, J., and V. V. Kapitonov, 2001 PIFs meet Tourists and Harbingers: a superfamily reunion. Proceedings of the National Academy of Sciences of the United States of America 98: 12315–12316.

Kalyaanamoorthy, S., B. Q. Minh, T. K. F. Wong, A. von Haeseler and L. S. Jermiin, 2017 ModelFinder: fast model selection for accurate phylogenetic estimates. Nature Methods 14: 587–589.

Kanzaki, N., I. J. Tsai, R. Tanaka, V. L. Hunt, D. Liu et al., 2018 Biology and genome of a newly discovered sibling species of Caenorhabditis elegans. Nature Communications 9: 3216–3215.

Kapitonov, V. V., and J. Jurka, 1999 Molecular paleontology of transposable elements from Arabidopsis thaliana. Genetica 107: 27–37.

Katoh, K., and D. M. Standley, 2013 MAFFT multiple sequence alignment software version 7: improvements in performance and usability. Molecular Biology and Evolution 30: 772–780.

Kawahara, K., T. Inada, R. Tanaka, M. Dayi, T. Makino et al., 2023 Differentially Expressed Genes Associated with Body Size Changes and Transposable Element Insertions between Caenorhabditis elegans and Its Sister Species, Caenorhabditis inopinata. Genome Biology and Evolution 15: evad063.

Ko, F. C. F., and K. L. Chow, 2002 A novel thioredoxin-like protein encoded by the C. elegans dpy-11 gene is required for body and sensory organ morphogenesis. Development (Cambridge, England) 129: 1185–1194.

Liu, P., K. Panda, S. A. Edwards, R. Swanson, H. Yi et al., 2024 Transposase-assisted target-site integration for efficient plant genome engineering. Nature 631: 593–600.

Minh, B. Q., H. A. Schmidt, O. Chernomor, D. Schrempf, M. D. Woodhams et al., 2020 IQ-TREE 2: New Models and Efficient Methods for Phylogenetic Inference in the Genomic Era. Molecular Biology and Evolution 37: 1530–1534.

Mistry, J., S. Chuguransky, L. Williams, M. Qureshi, G. A. Salazar et al., 2021 Pfam: The protein families database in 2021. Nucleic Acids Research 49: D412–D419.

Oomura, S., K. Tsuyama, N. Haruta and A. Sugimoto, 2022 Transgenesis of the gonochoristic nematode Caenorhabditis inopinata by microparticle bombardment with hygromycin B selection. microPublication Biology 2022: 10.17912/micropub.biology.000564. eCollection 002022.

Rice, P., I. Longden and A. Bleasby, 2000 EMBOSS: the European Molecular Biology Open Software Suite. Trends in Genetics : TIG JID507085 16: 276–277.

Rognes, T., T. Flouri, B. Nichols, C. Quince and F. Mahe, 2016 VSEARCH: a versatile open source tool for metagenomics. PeerJ 4: e2584.

Rubin, E., and A. A. Levy, 1997 Abortive gap repair: underlying mechanism for Ds element formation. Molecular and Cellular Biology 17: 6294–6302.

Shen, W., S. Le, Y. Li and F. Hu, 2016 SeqKit: A Cross-Platform and Ultrafast Toolkit for FASTA/Q File Manipulation. PLoS One 11: e0163962.

Sinzelle, L., V. V. Kapitonov, D. P. Grzela, T. Jursch, J. Jurka et al., 2008 Transposition of a reconstructed Harbinger element in human cells and functional homology with two transposon-derived cellular genes. Proceedings of the National Academy of Sciences of the United States of America 105: 4715–4720.

Siomi, M. C., K. Sato, D. Pezic and A. A. Aravin, 2011 PIWI-interacting small RNAs: the vanguard of genome defence. Nature Reviews.Molecular Cell Biology 12: 246–258.

Sun, S., N. Kanzaki, M. Dayi, Y. Maeda, A. Yoshida et al., 2022 The compact genome of Caenorhabditis niphades n. sp., isolated from a wood-boring weevil, Niphades variegatus. BMC Genomics 23: 765–768.

Velanis, C. N., P. Perera, B. Thomson, E. de Leau, S. C. Liang et al., 2020 The domesticated transposase ALP2 mediates formation of a novel Polycomb protein complex by direct interaction with MSI1, a core subunit of Polycomb Repressive Complex 2 (PRC2). PLoS Genetics 16: e1008681.

Vora, M., J. Dietz, Z. Wing, K. George, J. Kelly Liu et al., 2025 Genome-wide analysis of Smad and Schnurri transcription factors in C. elegans demonstrates widespread interaction and a function in collagen secretion. eLife 13: 10.7554/eLife.99394.

Wang, K., M. Li and H. Hakonarson, 2010 ANNOVAR: functional annotation of genetic variants from high-throughput sequencing data. Nucleic Acids Research 38: e164.

Wicker, T., F. Sabot, A. Hua-Van, J. L. Bennetzen, P. Capy et al., 2007 A unified classification system for eukaryotic transposable elements. Nature Reviews.Genetics 8: 973–982.

Yang, G., D. H. Nagel, C. Feschotte, C. N. Hancock and S. R. Wessler, 2009 Tuned for transposition: molecular determinants underlying the hyperactivity of a Stowaway MITE. Science (New York, N.Y.) 325: 1391–1394.

Yang, G., F. Zhang, C. N. Hancock and S. R. Wessler, 2007 Transposition of the rice miniature inverted repeat transposable element mPing in Arabidopsis thaliana. Proceedings of the National Academy of Sciences of the United States of America 104: 10962–10967.

Zhang, X., C. Feschotte, Q. Zhang, N. Jiang, W. B. Eggleston et al., 2001 P instability factor: an active maize transposon system associated with the amplification of Tourist-like MITEs and a new superfamily of transposases. Proceedings of the National Academy of Sciences of the United States of America 98: 12572–12577.

Zhou, L., T. Feng, S. Xu, F. Gao, T. T. Lam et al., 2022 ggmsa: a visual exploration tool for multiple sequence alignment and associated data. Briefings in Bioinformatics 23.

